# Demographic causes of adult sex ratio variation and their consequences for parental cooperation

**DOI:** 10.1101/223941

**Authors:** Luke J. Eberhart-Phillips, Clemens Küpper, María Cristina Carmona-Isunza, Orsolya Vincze, Sama Zefania, Medardo Cruz-López, András Kosztolányi, Tom E. X. Miller, Zoltán Barta, Innes C. Cuthill, Terry Burke, Tamás Székely, Joseph I. Hoffman, Oliver Krüger

**Author notes:** Joint senior authors (Tamás Székely, Joseph I. Hoffman, Oliver Krüger). **Correspondence to:** Luke J. Eberhart-Phillips, Research Group Behavioural Genetics and Evolutionary Ecology Max Planck Institute for Ornithology Eberhard-Gwinner-Str. 5, 82319 Seewiesen, Germany.

## Abstract

The adult sex ratio (ASR) is a fundamental concept in population biology, sexual selection, and social evolution. However, it remains unclear which demographic processes generate ASR variation and how biases in ASR in turn affect social behaviour. Here, we evaluate the demographic mechanisms shaping ASR and their consequences for parental cooperation using detailed survival, fecundity, and behavioural data on 6,119 individuals from six wild shorebird populations exhibiting flexible parental strategies. We show that these closely related populations express strikingly different ASRs, despite having similar ecologies and life histories, and that ASR variation is largely driven by sex differences in the apparent survival of juveniles. Furthermore, families in populations with biased ASRs were predominantly tended by a single parent, suggesting that parental cooperation breaks down with unbalanced sex ratios. Taken together, our results indicate that sex biases emerging during early life have profound consequences for social behaviour.

## Main text

Sex ratio variation is a fundamental component of life-history evolution. At conception, birth, and adulthood, the ratios of males to females have long been hypothesized by evolutionary biologists and human demographers as catalysts for social behaviour and population dynamics^1,2^. In particular, the adult sex ratio (ASR) exhibits remarkable variation throughout nature, with birds and mammals tending to have male-biased and female-biased ASRs, respectively^3^. Recent studies also show extreme shifts in ASR due to climate change in fish^4^, amphibians^5^, and dioecious plants^6^. By influencing mate availability, ASR bias can alter social behaviour with divorce, infidelity, and parental antagonism being more frequent in sex-biased populations^7,8^. Moreover, in human societies, ASR variation is linked to economic decisions, community violence, and disease prevalence^9-11^. Yet despite the widespread occurrence of ASR bias and its significance in evolutionary ecology and social science, the demographic source(s) of ASR variation and their ramifications for social behaviour remain unclear^12^.

Sex ratio theory is concerned with the adaptive consequences of sex-biased parental allocation to offspring^13,14^, with the processes generating sex ratio bias after birth receiving less theoretical and empirical attention^15^. Here, we use a demographic pathway model to quantify ASR variation among avian populations and to determine whether this variation is predominantly caused by sex biases at birth, during juvenile development, or in adulthood. We parameterized our model with detailed individual-based life history data from *Charadrius* plovers – small ground-nesting shorebirds that occur worldwide. Plovers exhibit remarkable diversity and plasticity in breeding behaviour with sex roles during courtship, mating, and parental care varying appreciably among populations both between and within species^16,17^. This behavioural variation, coupled with their tractability in the field (Supplementary Video 1), allowed us to explore the sources and significance of demographic sex biases among closely related wild populations in the light of social evolution.

## Results and Discussion

Over a total of 43 observational years of fieldwork, we monitored the survival, fecundity, and breeding behaviour of 6,119 individually marked plovers from six populations of five closely related species worldwide (Fig. 1a). We then employed two-sex stage-structured population matrix models to derive estimates of ASR at equilibrium from stage- and sex-specific demographic rates of annual survival and reproduction (Fig. 1b)^18^. For each population, the numbers of male and female progeny in our model depended on modal clutch size and hatching sex ratio derived from our field data. Mark-recapture methods were used to estimate the apparent survival of juveniles and adults while accounting for sex differences in detection probability (the term “apparent survival” indicates that mortality cannot be disentangled from permanent emigration)^19^. Fecundity was derived from a mating function that depended on the extent of polygamy observed in each population and the frequency of available mates (see *Methods* for details).

**Figure 1.**
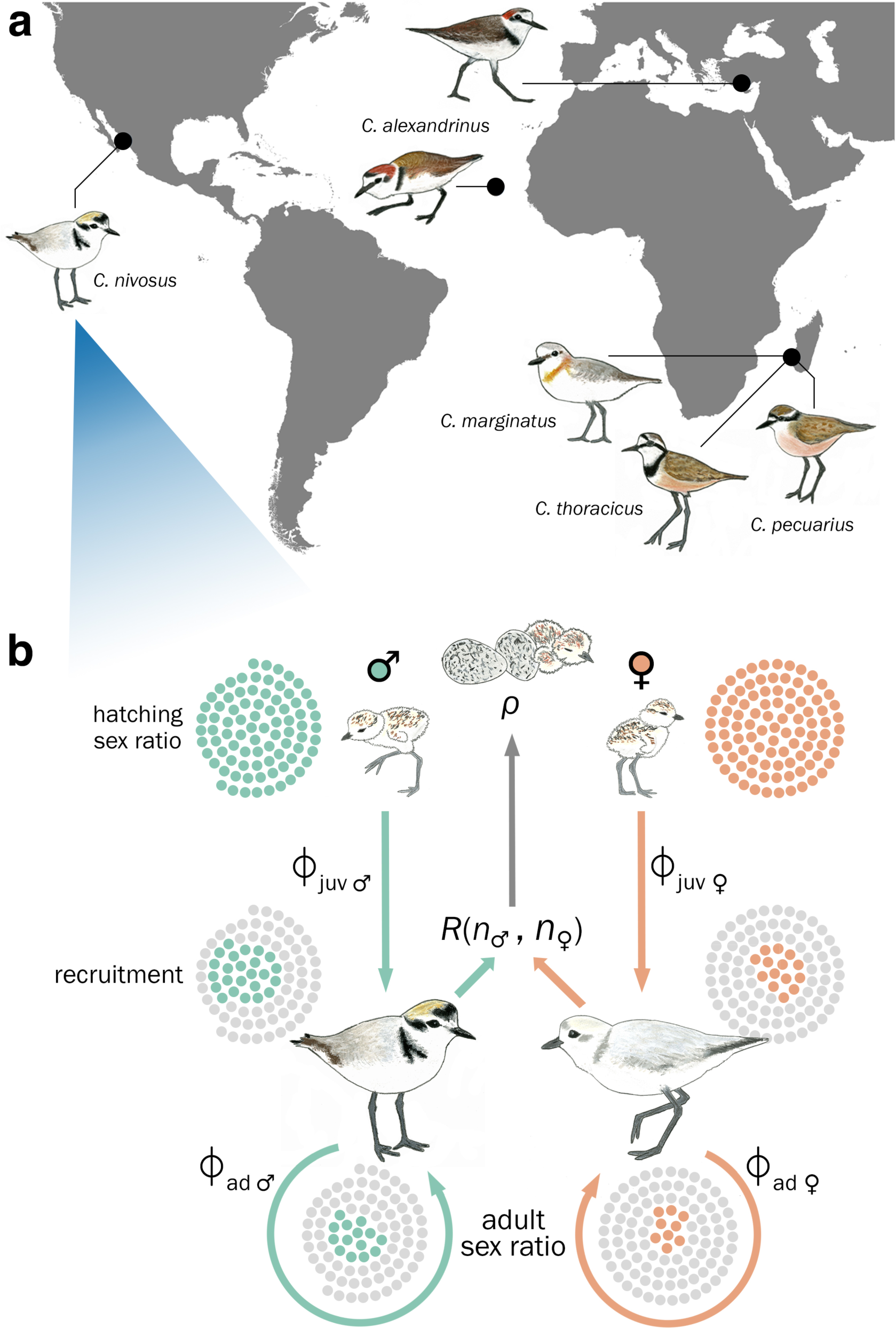
Modelling the demographic pathway of adult sex ratio bias among plovers worldwide. (**a**) Location of the six study populations. *C. pecuarius, C. marginatus,* and *C. thoracicus* breed sympatrically in south-western Madagascar, whereas the two populations of *C. alexandrinus* are geographically disparate, inhabiting southern Turkey and the Cape Verde archipelago. The studied *C. nivosus* population is located on the Pacific coast of Mexico. All populations inhabit saltmarsh or seashore habitats characterized by open and flat substrates. (**b**) Schematic of the stage- and sex-specific demographic transitions of individuals from hatching until adulthood and their contributions to the adult sex ratio (depicted here is *C. nivosus).* The hatching sex ratio (*ρ*, proportion of male hatchlings) serves as a proxy for the primary sex ratio and allocates progeny to the male or female juvenile stage. During the juvenile (‘juv’) stage, a subset of this progeny will survive (ϕ) to recruit and remain as adults (‘ad’). Dotted clusters illustrate how a cohort is shaped through these sex-specific demographic transitions to derive the adult sex ratio (mortality indicated by grey dots). The reproduction function, *R*(*n*_♂_, *n*_♀_), is dependent on mating system and the frequency of available mates (see *Methods* for details).

The hatching sex ratio, based on 1,139 hatchlings from 503 families, did not deviate significantly from parity in any of the populations (Fig. 2a). Conversely, sex biases in apparent survival varied considerably within and among species and, in most populations, juvenile survival was more biased than adult survival – either towards males or females (Fig. 2a). Taken together, these sources of demographic sex bias rendered significant deviations in ASR from parity for three populations (two male-biased and one female-biased; Fig. 2b).

**Figure 2.**
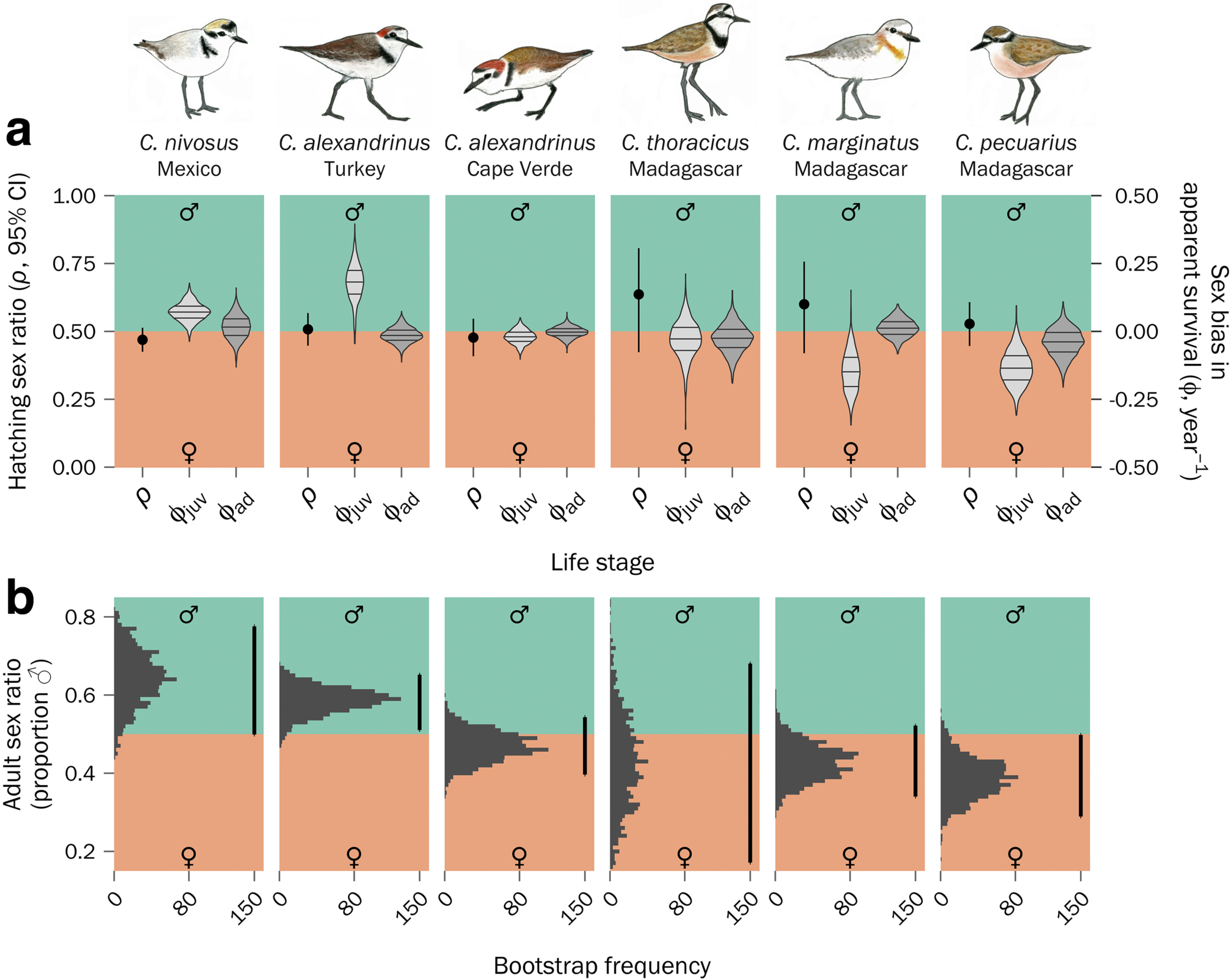
Inter- and intra-specific variation in sex-biased demography. (**a**) Hatching sex ratios of successful clutches (proportion of chicks that are male) are shown as point estimates (*ρ* ± 95% CI; left y-axis), and sex bias (i.e. difference between males and females) in annual apparent survival rates of juveniles (ϕ_juv_) and adults (ϕ_ad_) are shown as violin plots (right y-axis). Horizontal lines within violin plots indicate the median and interquartile ranges of the bootstrapped estimates (see *Methods* for details). (**b**) Bootstrap distributions of the derived ASRs based on the sex- and stage-specific rates shown in panel a. Vertical bars on the right side of histograms indicate the 95% CI of ASRs based on 1,000 iterations of the bootstrap (mean ASR [95% CI]: *C. nivosus* = 0.644 [0.499, 0.778], *C. alexandrinus* [Turkey] = 0.586 [0.510, 0.652], *C. alexandrinus* [Cape Verde] = 0.469 [0.396, 0.543], *C. thoracicus* = 0.421 [0.171, 0.681], *C. marginatus* = 0.430 [0.340, 0.522], *C. pecuarius* = 0.386 [0.289, 0.498].

Matrix models provide a flexible analytical environment to decompose the feedbacks between state-dependent vital rates and population response – an important method used in conservation biology for understanding life-history contributions to population growth and viability^20^. In our case, we modified this approach to assess the relative contributions of sex allocation and sex-specific survival on ASR bias^18^. We found that sex biases in apparent survival during the juvenile stage contributed the most to sex ratio bias of the adult population: in populations with significantly skewed ASR, sex biases in juvenile survival contributed on average 7.2 times more than sex biases in adult apparent survival and 24.6 times more than sex biases at hatching (Supplementary Fig. 1). Moreover, variation in hatching sex ratio had no effect on ASR – a result that provides empirical support for Fisher’s^13^ prediction of unbiased sex allocation regardless of sex-biased survival of independent young or adults.

In species where both parents have equal caring capabilities, the desertion of either parent is often influenced by the availability of potential mates^21^ – parental care by the abundant sex is expected to be greater than that of the scarcer sex due to limited future reproductive potential^22^. Detailed behavioural observations of 471 plover families revealed high rates of parental desertion in populations with biased ASRs, whereas desertion was rare in unbiased populations (Fig. 3). We evaluated our *a priori* prediction of a quadratic relationship between parental cooperation and ASR variation using a regression analysis incorporating a bootstrap procedure that acknowledged uncertainty in our estimates of ASR and parental care (see *Methods* for details). We found that families in male- or female-biased populations tended to express higher rates of parental desertion, while unbiased populations were more likely to exhibit parental cooperation (Fig. 3a). This is supported by experimental evidence of sex-biased mating opportunities in three of the populations studied here (Supplementary Fig. 2; ref 23). Moreover, the relationship between parental cooperation and local ASR bias was apparent in our within-species contrast of *C. alexandrinus:* the unbiased Cape Verde population exhibited a higher rate of parental cooperation than the male-biased population in Turkey (Fig. 3a). Counterintuitively, we also found a high rate of male-only care in *C. pecuarius* despite ASR being female-biased (Fig. 3b), although in line with expectations, *C. pecuarius* also showed the highest proportion of female-only care among our studied populations (Fig. 3b, Supplementary Fig. 3b). This provides partial support for the notion that breeding strategies may respond flexibly to local mating opportunities provided by ASR bias, while also suggesting that other factors may play a role, such as the energetic costs of egg production imposed on females or because of sex differences in parental quality^24^ or the age at maturation^25^.

**Figure 3.**
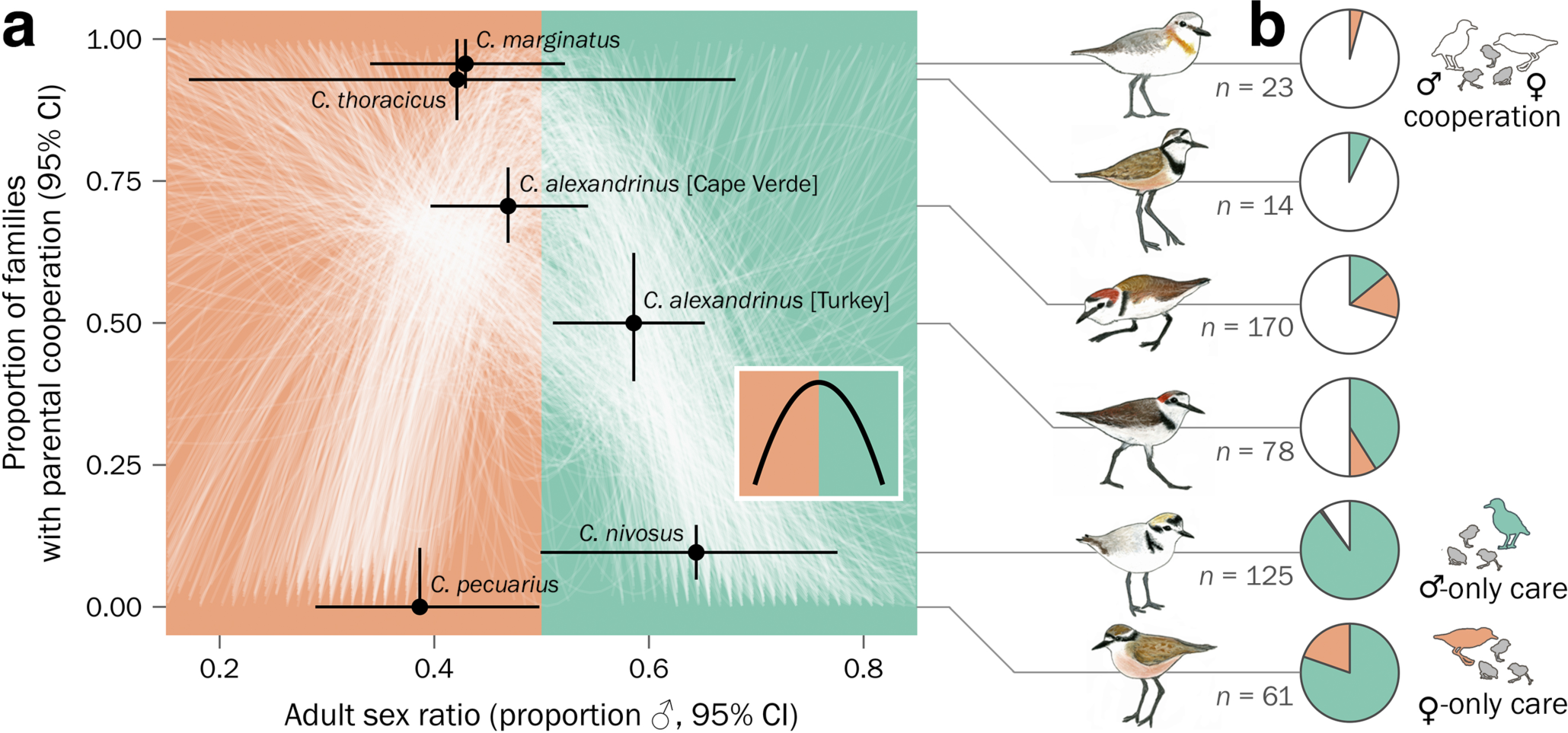
Relationship between parental cooperation and the adult sex ratio. (**a**) Faint white lines illustrate each iteration of the bootstrap, which randomly sampled an adult sex ratio and parental care estimate from each population’s uncertainty distribution and fitted them to the *a priori* quadratic model (shown in inset, Eq. 11). (**b**) Proportion of monitored plover families that exhibit parental cooperation (white) or single-parent care by males (green) or females (orange). Sample sizes reflect number of families monitored per population.

There are several ways in which sex-biased juvenile survival could potentially lead to a skewed ASR. Natal dispersal rates may differ between the sexes, as is typical of many birds^26^, which could contribute to our estimates of sex-biased apparent survival. Genetic studies of several of the populations presented here are partially consistent with this hypothesis, as island populations of *C. alexandrinus* and the endemic *C. thoracicus* have reduced gene flow relative to comparable mainland populations^27,28^. However, sex-biased juvenile survival in plovers has been reported elsewhere, even after accounting for dispersal^29,30^, implying that sex-differences in mortality are at least partly driven by intrinsic factors, conceivably for example via genotype-sex interactions^31,32^. An alternative but not necessarily mutually exclusive explanation is that males and females may differ in their premature investment into reproductive traits, which could inflict survival costs for the larger or more ornamented sex^33^. Although sexual dimorphism among adults is negligible in the species we studied^34,35^, sex-specific ontogeny does appear to vary in three of these populations. In male-biased populations of *C. nivosus* and *C. alexandrinus,* female hatchlings are smaller and grow more slowly than their brothers during the first weeks of life, whereas juveniles of the unbiased *C. alexandrinus* population exhibit no such sex-specific differences during early development^34^.

The association between sex-specific demography and breeding system evolution represents a causality dilemma because of the feedback that parental strategies impose on ASR bias and *vice versa*^36^. On the one hand, mating competition and parental care may entail costs via sexual selection that could drive differential survival of males over females and have knock-on effects on ASR^37^. On the other hand, sex-specific survival creates unequal mating opportunities via ASR that may influence mating patterns and parenting strategies^37^. Our study provides empirical support for the latter – sex biases emerge prior to sexual maturity, suggesting that this evolutionary feedback loop is catalysed by intrinsic early-life demographic variation. Moreover, our results add to the growing evidence of unbiased birth sex ratios in nature^15^ and provide a comprehensive empirical test of Fisher’s^13^ original prediction that influential sex biases arise in life-history stages beyond parental control. By unravelling the demographic foundations of adult sex ratio bias and their consequences for parental cooperation, we hope to stimulate future studies to understand the complex relationship between evolutionary demography and behavioural ecology.

## Methods

### Field and laboratory methods

We studied five *Charadrius* species comprising six populations from four sites worldwide (Fig. 1a, Supplementary Table 1). In Mexico, we monitored the snowy plover (*C*. *nivosus*) at Bahía de Ceuta, a subtropical lagoon on the Pacific coast. In Madagascar, we monitored the Kittlitz’s plover (*C*. *pecuarius*), white-fronted plover (*C*. *marginatus*), and the endemic Madagascar plover (*C*. *thoracicus*), all of which breed sympatrically at a saltwater marsh near the fishing village of Andavadoaka. Lastly, we monitored the Kentish plover (*C*. *alexandrinus*) at two independent populations located at Lake Tuzla in southern Turkey and at Maio in Cape Verde. The Mexico and Madagascar populations were monitored over a seven-year period, whereas the Turkey and Cape Verde populations were monitored over six and nine years respectively, thus totalling 43 years of data collection (Supplementary Table 1).

At each location, we collected mark-recapture and individual reproductive success data during daily surveys over the entire breeding season that typically spanned three to four months after a region’s rainy season. Funnel traps were used to capture adults on broods or nests^38^. We assigned individuals to a unique colour combination of darvic rings and an alphanumeric metal ring, allowing the use of both captures and non-invasive resightings to estimate survival. Broods were monitored every five days on average to assess daily survival and identify tending parents. During captures, approximately 25–50 μL of blood was sampled from the metatarsal vein of chicks and the brachial vein of adults for molecular sex typing with two independent markers Z-002B^39^ and Calex-31, a microsatellite marker on the W chromosome^40^. Details of PCR conditions are given in ref. 34.

### Quantifying sex roles

We evaluated sex role variation by summarizing for each population the proportion of all families that were attended bi-parentally or uni-parentally by a male or female. We restricted our field observations to include only broods that were at least 20 days old. Young chicks are attended by both parents in all populations although, as broods get older, male or female parents may desert the family^41^. As chicks typically fledge around 25 days of age, we therefore choose broods of between 20 and 25 days of age to quantify parental care given that at this age many parents already deserted the family but some still attend the young. Furthermore, we restricted these data to include only broods that had at least two sightings after 20 days. Given these criteria, our dataset consisted of 471 unique families distributed throughout the six populations and pooled across all years of study (Supplementary Table 2). To account for surveyor oversight while recording tending parents (e.g., observing only one parent when two were present), we took a conservative approach by assigning a bi-parental status to families that had both uni-parental and bi-parental observations after the 20^th^ day. In summary, desertion was most common in *C. nivosus* and *C. pecuarius*, whereas bi-parental care was most common in *C. alexandrinus* (Cape Verde), *C. thoracicus*, and *C. marginatus* (Fig. 3b, Supplementary Table 2). The *C. alexandrinus* population in Turkey had 50% bi-parental and 50% desertion (Fig. 3b, Supplementary Table 2). *C. pecuarius* had the highest incidence of male desertion (20%; Fig. 3b, Supplementary Table 2). To acknowledge uncertainty in these parental care proportions given variation in sample size, we used a method that estimated simultaneous 95% confidence intervals according to a multinomial distribution^42^ (Supplementary Table 2).

### Estimation of sex- and stage-specific survival

Our structured population model considered sex-specific survival during two key stage classes in life history: juveniles and adults (Fig. 1b). The juvenile stage was defined as the one-year transition period between hatching and recruitment into the adult population. The adult stage represented a stasis stage in which individuals were annually retained in the population.

We used mark-recapture models to account for sex, stage, and temporal variation in encounter (*p*) and apparent survival (ϕ) probabilities as they allow for imperfect detection of marked individuals during surveys and the inclusion of individuals with unknown fates^40^. We use the term “apparent survival” as true mortality cannot be disentangled from permanent emigration in this framework^19^. We used Cormack-Jolly-Seber models to estimate juvenile and adult survival, with one-year encounter intervals. Juvenile and adult survival models were constructed from design matrices that included sex, year, and stage as factors.

Since we were primarily interested in stage- and sex-specific variation in survival, all models included a ϕ ~ *sex* * *stage* component. Our model selection thus evaluated the best structure explaining variation in detection probability by comparing all interactions between sex, year, and stage (e.g., *p* ~ *sex* * *year* * *stage*). We constructed survival models with the R package “RMark”^43^ and estimated demographic parameters via maximum likelihood implemented in program MARK^44^. We evaluated whether our data were appropriately dispersed (i.e. *c-hat* ≤ 3; ref. 19) by employing the “median c-hat” goodness-of-fit bootstrap simulation in program MARK^44^.

### Estimating hatching sex ratios

To account for potential sex-biases arising prior to the juvenile stage (i.e. sex allocation or sex-specific embryo mortality), we tested whether the hatching sex ratio deviated significantly from parity. Each population was analysed separately using a general linear mixed effect model with a binomial error distribution and a logit link function (R package “lme4”^45^). In this model, the response variable was chick sex, the fixed effect was the intercept, and brood identifier was included as a random factor to control for the non independence of siblings from the same nest. Because plover chicks are precocial, post-hatch brood mixing can occur. Consequently, our dataset for analysing hatching sex ratio included only complete broods (i.e. with no missing chicks) that were captured at the nest on the same day of hatching (503 unique families with 1139 chicks, Supplementary Table 3). The fixed-effect intercepts of all populations were not significantly different from zero, indicating that hatching sex ratios did not deviate from parity (Fig. 2a).

### Matrix model structure

We built two-sex post-breeding matrix models for each plover population that incorporated two annual transitions denoting juveniles and adults (Fig. 1b). The projection of the matrix for one annual time step (*t*) is given by:

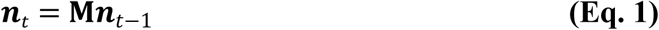

where *n* is a 4 × 1 vector of the population distributed across the two life stages and two sexes:

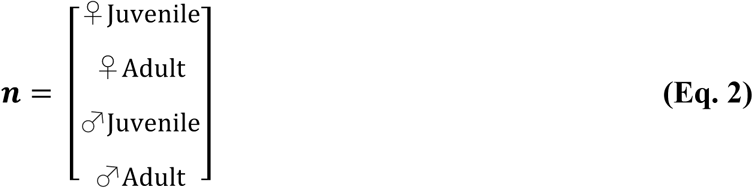

and **M** is expressed as a 4 × 4 matrix:

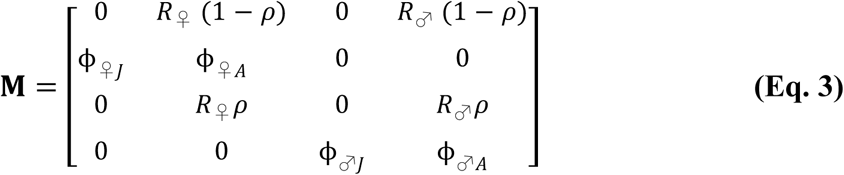

where transition probabilities (ϕ) between life stages are the apparent survival rates of female (♀) and male (♂) juveniles (*J*) and adults (*A*). The hatching sex ratio (*ρ*) describes the probability of hatchlings being either male (i.e. *ρ*) or female (i.e. 1 − *ρ*), and was estimated for each population from our field data (see above). *Per capita* reproduction of females (*R*_♀_) and males (*R*♂) is expressed through sex-specific mating functions used to link the sexes and produce progeny for the following time step of the model given the relative frequencies of each sex^20^. We used the harmonic mean mating function which accounts for sex-specific frequency dependence^46^:

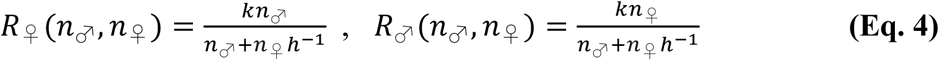

where *k* is the modal clutch size (3 in *C. nivosus, C. alexandrinus,* and *C. marginatus,* and 2 in *C. thoracicus* and *C. pecuarius*), *h* is an index of the annual number of mates acquired per female (i.e. mating system, see below), and *n*_♀_ and *n*_♂_ are the densities of females and males, respectively, in each time step of the model.

### Quantifying mating system

Demographic mating functions are traditionally expressed from the perspective of males^46^, whereby *h* is the average harem size (number of female mates per male). Under this definition, *h* > 1 signifies polygyny, *h* = 1 monogamy, and *h* < 1 polyandry; ref. 47). Although both sexes can acquire multiple mates in a single breeding season, within-season polygamy is typically female biased in plovers. Thus, in accordance with the predominantly polyandrous or monogamous mating systems seen across these six populations, *h* was derived from the average annual number of mates per female (*μ*):

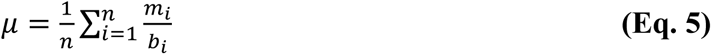

where, *n* is the total number of females in a given population, *b* is the total number of years female *i* was seen breeding, and *m* is the total number of mating partners female *i* had over *b* years. Thus, if *μ* was less than or equal to one, females tended to have only one mating partner annually, and *h* was set to 1. Alternatively, if *μ* was greater than 1, females were polygamous and *h* was calculated as the inverse of *μ*:

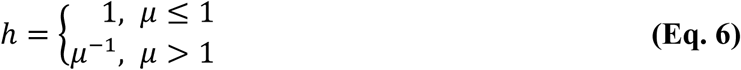

Our dataset to estimate *h* for each population only included females for which we were confident of the identity of their mates, and had observed them in at least two reproductive attempts. In summary, *μ* varied among populations (Supplementary Fig. 4), with *C. nivosus* (*h* = 0.82), *C. alexandrinus* (Turkey; *h* = 0.85), and *C. pecuarius* (*h* = 0.86) having polyandrous mating systems and *C. alexandrinus* (Cape Verde), *C. thoracicus,* and *C. marginatus* all having monogamous mating systems (i.e. *h* = 1).

### Estimation of the adult sex ratio

We estimated ASR from the stable stage distribution (**w**) of the two-sex matrix model:

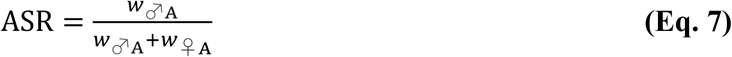

where *w*_♂A_ and *w*_♀A_ provide the proportion of the population composed of adult males and females, respectively, at equilibrium. To evaluate uncertainty in our estimate of ASR due to sampling and process variation in our apparent survival parameters, we implemented a bootstrapping procedure in which each iteration: (i) randomly sampled our mark-recapture data with replacement, (ii) ran the survival analyses described above, (iii) derived stage- and sex-specific estimates of apparent survival based on the model with the lowest AICc (i.e. ΔAICc = 0; Supplementary Fig. 5), (iv) constructed the matrix model (Eq. 3) of these estimates, (v) derived the stable stage distribution through simulation of 1,000 time steps, then (vi) derived ASR from the stable stage distribution at equilibrium on the 1,000^th^ time step. This approach ensured that parameter correlations within the matrix were retained for each bootstrap and it also accounted for non-linearity in the mating function. We ran 1,000 iterations and evaluated the accuracy of our ASR estimate by determining the 95% confidence interval of its bootstrapped distribution. Note that our method estimated ASR as the asymptotic value predicted under the assumption that each population was at equilibrium and thus we could not evaluate inter-annual variation in asymptotic ASR. Nonetheless, our model-derived ASR estimate of the *C. nivosus* population falls within annual count-based ASR estimates of this population^48^, providing support that our method is robust. Count-based estimates of ASR from the remaining populations in our study are unfortunately uninformative due to our limited sample of marked individuals with known sex.

Our mark-recapture analysis was based on the encounter histories of 6,119 uniquely marked and molecularly sexed individuals (Supplementary Table 4). After implementing the bootstrap procedure, we found that variation in the encounter probabilities of juveniles and adults was best explained by sex, year, and age in *C. nivosus*, *C. alexandrinus* (Turkey) and *C. pecuarius* (Supplementary Fig. 5). Encounter probability was best explained by age and year in *C. alexandrinus* (Cape Verde) and *C. marginatus* (Supplementary Fig. 5). In *C. thoracicus,* encounter probability was best explained by sex and year (Supplementary Fig. 5). Our mark-recapture data were not over-dispersed (Supplementary Table 4).

### Life table response experiment of ASR contributions

Perturbation analyses provide information about the relative effect that each component of a matrix model has on the population-level response, in our case ASR. To assess how influential sex biases in parameters associated with each of the three life stages were on ASR dynamics, we employed a life-table response experiment (LTRE). A LTRE decomposes the difference in response between two or more “treatments” by weighting the difference in parameter values by the parameter's contribution to the response (i.e. its sensitivity), and summing over all parameters^20^. We compared the observed scenario (**M**), to a hypothetical scenario (**M**_0_) whereby all female survival rates were set equal to the male rates (or *vice versa)* and the hatching sex ratio was unbiased (i.e. *ρ* = 0.5). Thus, our LTRE identifies the drivers of ASR bias by decomposing the difference between the ASR predicted by our model and an unbiased ASR^18^.

The contributions (C) of lower-level demographic parameters (θ) were calculated following Veran and Beissinger^18^:

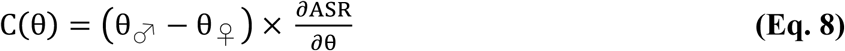

where 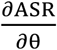 is the sensitivity of ASR to perturbations in the demographic rate θ in matrix **M**’, which is a reference matrix “midway” between the two scenarios^18^:

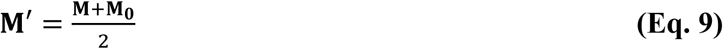

The two-sex mating function makes our model non-linear in the sense that the projection matrix, and specifically the fertility elements (Eq. 4), depends on sex-specific population structure. Perturbation analyses must therefore accommodate the indirect effects of parameter perturbations on population response via their effects on population structure, such as the relative abundance of males and females which can affect mating dynamics and fecundity. To estimate the sensitivities of the ASR to vital rate parameters, we employed numerical methods that independently perturbed each parameter of the matrix, simulated the model through 1,000 time steps, and calculated ASR at equilibrium. This produced parameter-specific splines from which 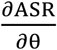 could be derived. This approach appropriately accounts for the non-linear feedbacks between vital rates and population structure, though it does not isolate the contribution of this feedback^47,49^.

Our LTRE revealed that across all populations, sex differences in juvenile apparent survival made the largest overall contribution to ASR bias (Supplementary Fig. 1). Likewise, for all populations, sex biases at hatching and in mating system had negligible effects on ASR variation (Supplementary Fig. 1).

### Evaluating the association between ASR bias and parental cooperation

To test the relationship between ASR bias and parental cooperation, we conducted a regression analysis of the following quadratic model:

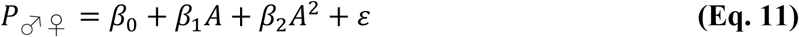

where ***P***_♂♀_ is the proportion of families exhibiting parental cooperation, *β_i_* are the regression parameters (i.e. intercept and coefficient), *A* is the ASR, and *ε* is random error. We chose a quadratic model *a priori* as we expected maximum parental cooperation at unbiased ASR but minimum cooperation at both male- and female-biased ASRs (see inset in Fig. 3a).This relationship was assessed with a bootstrap procedure that incorporated uncertainty in our estimates of ASR and parental care. Each iteration of the bootstrap (i) randomly sampled an ASR value from the 95% confidence interval of each population shown in Fig. 2b, (ii) randomly sampled a parental care value from the truncated 95% confidence interval of each population shown in Supplementary Table 2, then (iii) fitted the regression model. We ran 1,000 iterations of the bootstrap and evaluated overall relationships by visualizing the central tendency of the regressions. We also evaluated the relationship between ASR variation and male-only or female-only care using a similar bootstrap procedure of the following models:

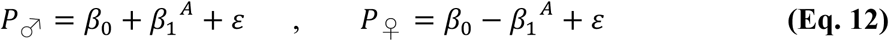

where *P*_♂_ and *P*_♀_ are the proportions of families exhibiting male-only or female-only care, respectively. In this case, we chose exponential models *a priori* as we expected a non-linear increase in uni-parental care by the abundant sex under biased ASR (Supplementary Fig. 3a). This analysis demonstrated that male-only care tended to be more common in populations with male-biased ASR (mean *β*_1_ = 0.682 [-0.366, 1.555 95% CI]) and female-only care tended to be more common in female-biased populations (mean *β*_1_ = -0.205 [-0.502, 0.037 95% CI]; Supplementary Fig. 3b). However, the overall magnitude of the effect of ASR variation on female-only care was less than that of male-only care.

### Data and code availability

All of our modelling and statistical analyses were conducted using R version *Kite-eating Tree*^50^ with significance testing evaluated at α = 0.05. We provide all computer code and documentation as a PDF file written in RMarkdown (Supplementary Material A) together with all the raw datasets needed to reproduce our modelling and analyses (Supplementary Material B). These can be downloaded from our GitHub repository: https://github.com/leberhartphillips/Plover_ASR_Matrix_Modeling.

## Acknowledgements

Fieldwork was permitted and ethically approved by federal authorities in Cape Verde (Direcção Geral do Ambiente, DGA), Mexico (Secretaría de Medio Ambiente y Recursos Naturales, SEMARNAT), Madagascar (Ministry of Environment, Forests, and Tourism of the Republic of Madagascar), and Turkey (Turkish Ministry of Environment). Financial support was provided by a Deutsche Forschungsgemeinschaft (DFG) doctoral fellowship to LE-P and supervised by OK, JH, and TS (GZ: KR 2089/9-1). Fieldwork was funded by Sonoran Joint Venture, Tracy Aviary Conservation Fund, Convocatoria de Investigación Científica Básica (CONACyT, Grant no. 157570), Fundação Maio Biodiversidade (FMB), DGA, Camara Municipal do Maio, DFG Mercator Visiting Professorship awarded to TS, Natural Environment Research Council grant (GR3/10957), Biotechnology and Biological Sciences Research Council (BBS/B/05788) to ICC and TS. MCC-I: CONACyT (PhD scholarship 216052/311485). OV: Hungarian Eötvös Scholarship (Tempus Public Foundation MÁEÖ2016_15/76740), Campus Mundi scholarship (CMP-69-2/2016), and Hungarian Research Fund (OTKA #K113108). AK: János Bolyai Research Scholarship of the Hungarian Academy of Sciences and NKFIH grant (K112670). TB: Leverhulme Fellowship. TS: NKFIH-2558-1/2015 and the Wissenschaftskollegium zu Berlin. Our manuscript benefitted from the constructive comments of S. Beissinger, W. Forstmeier, M. Galipaud, M. Jennions, P. Korsten, A. Shah, M. Stoffel, and three anonymous reviewers. Original plover illustrations by LE-P. We thank N. dos Remedios and K. Maher for molecular work and we are grateful for the generous assistance of countless students, volunteers, and colleagues who aided with fieldwork.

## Author contributions

L.E.-P., T.S., J.I.H., and O.K. conceived the study. L.E.-P., C.K., M.C.C.-I., O.V., S.Z., A.K., M.C.L, I.C.C., and T.S. planned and collected the field data. L.E.-P., C.K., J.I.H., and T.B. performed or supervised the molecular sexing. L.E.-P., T.M., and O.K. implemented the demographic modelling. L.E.-P. wrote the manuscript and the RMarkdown file. All authors contributed substantially to revisions of the paper.

## Supplementary Information

***Supplementary Tables***

**Supplementary Table 1.** Study site metadata: geographic location (see Fig. 1) and duration of monitoring effort.

| Species | Population | Latitude | Longitude | Years<br>monitored |
| --- | --- | --- | --- | --- |
| *C. nivosus* | Mexico | 23°54'N | 106°57'W | 2006-2012 |
| *C. alexandrinus* | Turkey | 36°43'N | 35°03'E | 1996-2001 |
|  | Cape Verde | 15°8'N | 23°13'W | 2007-2015 |
| *C. thoracicus* | Madagascar | 22°6'S | 43°15'E | 2009-2015 |
| *C. marginatus* | Madagascar |  |  | 2009-2015 |
| *C. pecuarius* | Madagascar |  |  | 2009-2015 |
|  |  |  |  | 43 years |

**Supplementary Table 2.**
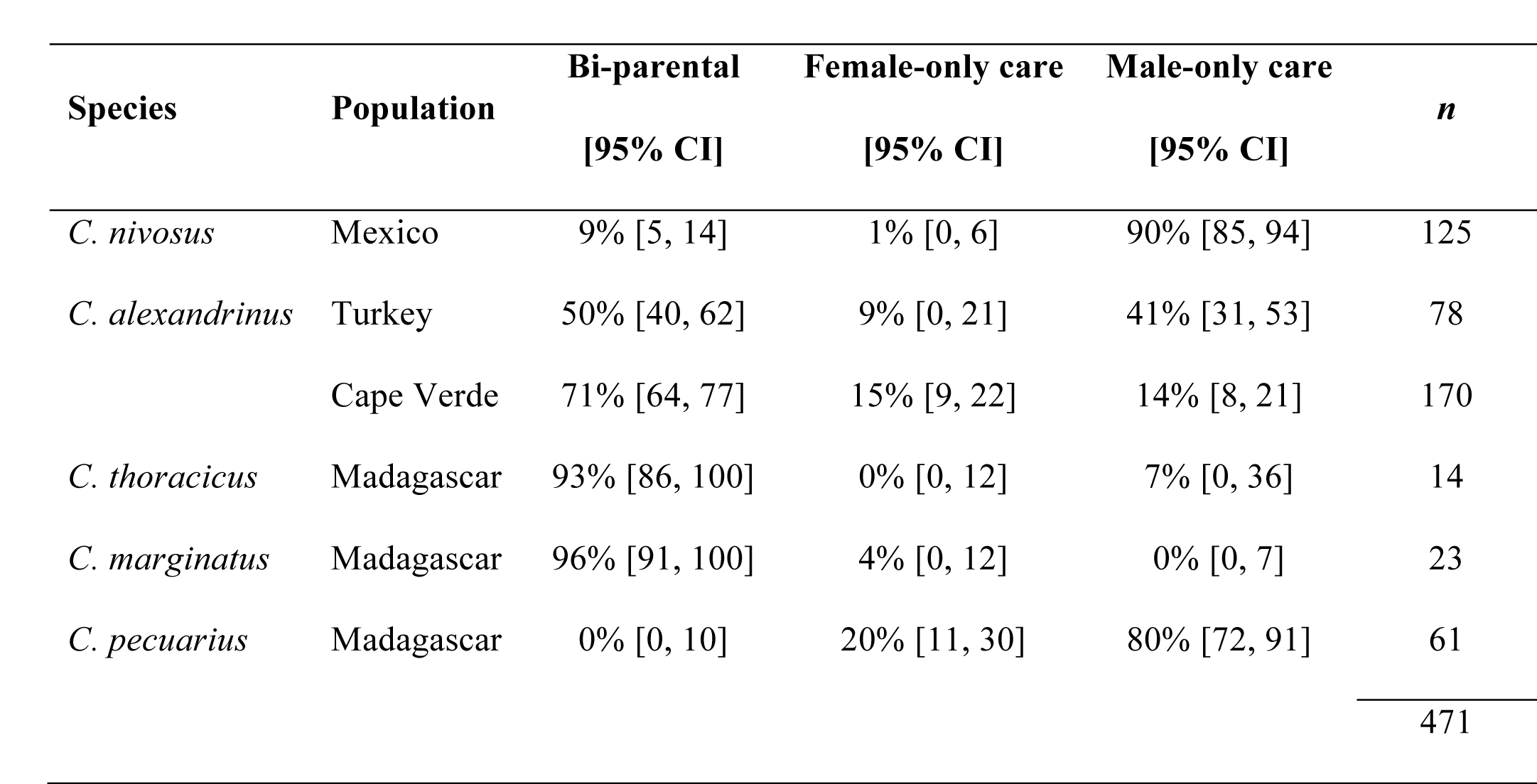
**Summary of parental care sex roles**. Percentages reflect the within-population proportion of families with a given parental care system.

| Species | Population | Bi-parental<br>[95% CI] | Female-only care<br>[95% CI] | Male-only care<br>[95% CI] | *n* |
| --- | --- | --- | --- | --- | --- |
| *C. nivosus* | Mexico | 9% [5, 14] | 1% [0, 6] | 90% [85, 94] | 125 |
| *C. alexandrinus* | Turkey | 50% [40, 62] | 9% [0, 21] | 41% [31, 53] | 78 |
|  | Cape Verde | 71% [64, 77] | 15% [9, 22] | 14% [8, 21] | 170 |
| *C. thoracicus* | Madagascar | 93% [86, 100] | 0% [0, 12] | 7% [0, 36] | 14 |
| *C. marginatus* | Madagascar | 96% [91, 100] | 4% [0, 12] | 0% [0, 7] | 23 |
| *C. pecuarius* | Madagascar | 0% [0, 10] | 20% [11, 30] | 80% [72, 91] | 61 |
|  |  |  |  |  | 471 |

**Supplementary Table 3.** **Summary of hatching sex ratio data**, where *ρ* is the average hatching sex ratio (expressed as the proportion of hatchlings in a brood that are male) and 95% CIs are calculated using a binomial distribution.

| Species | Population | $N_{\text{Families}}$ | $N_{\text{Hatchlings}}$ | $\rho$ | 95% CI |
| --- | --- | --- | --- | --- | --- |
| *C. nivosus* | Mexico | 198 | 484 | 0.469 | [0.425, 0.514] |
| *C. alexandrinus* | Turkey | 102 | 262 | 0.508 | [0.447, 0.568] |
|  | Cape Verde | 107 | 197 | 0.477 | [0.408, 0.547] |
| *C. thoracicus* | Madagascar | 11 | 22 | 0.636 | [0.423, 0.807] |
| *C. marginatus* | Madagascar | 13 | 30 | 0.600 | [0.419, 0.757] |
| *C. pecuarius* | Madagascar | 72 | 144 | 0.528 | [0.446, 0.757] |

**Supplementary Table 4.** Sample size and over-dispersion summary of mark-recapture dataset used to estimate apparent survival.

| Species | Population | Juveniles<sup>1</sup> | | Adults<sup>2</sup> | | Total | Median $\hat{c}$ |
| --- | --- | --- | --- | --- | --- | --- | --- |
|  |  | ♀ | ♂ | ♀ | ♂ | individuals |  |
| *C. nivosus* | Mexico | 438 | 388 | 221 | 212 | 1358 | 1.70 |
| *C. alexandrinus* | Turkey | 310 | 293 | 557 | 504 | 1664 | 1.49 |
|  | Cape Verde | 377 | 383 | 254 | 213 | 1227 | 1.37 |
| *C. thoracicus* | Madagascar | 38 | 56 | 83 | 68 | 245 | 2.72 |
| *C. marginatus* | Madagascar | 76 | 96 | 99 | 95 | 366 | 1.31 |
| *C. pecuarius* | Madagascar | 274 | 286 | 382 | 416 | 1358 | 1.77 |
|  |  |  |  |  |  | 6119 |  |

<sup>1</sup>Individuals first marked as hatchlings (i.e. known age).
<sup>2</sup>Individuals first marked as breeding adults (i.e. 1+ years old).

**Supplementary Figure 1.**
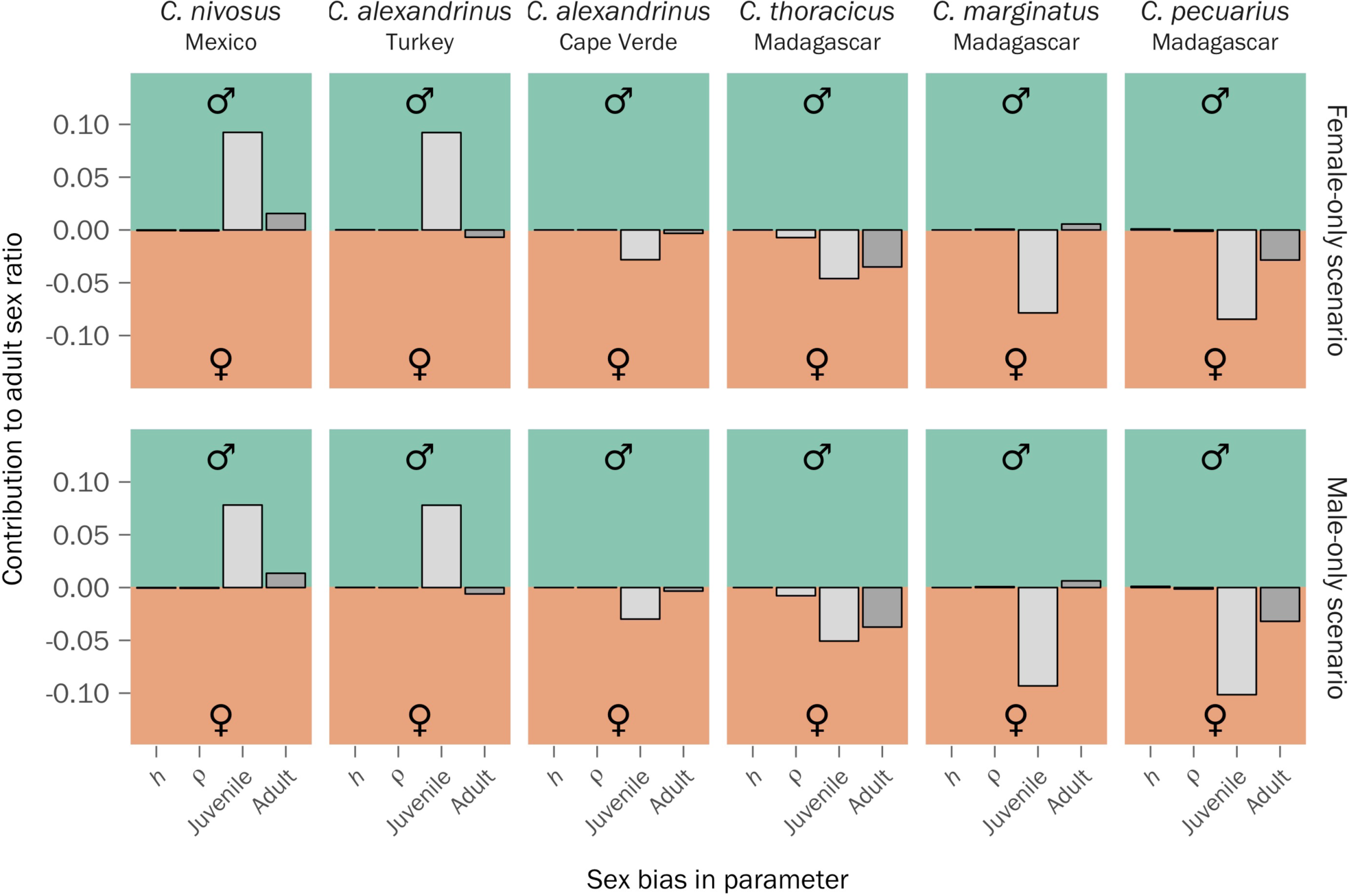
Contributions of sex-specific parameters to adult sex ratio bias. These results are based on a life-table response experiment (LTRE) that compared the empirically-derived sex-specific model to hypothetical scenarios with no sex differences in demographic rates (top panel: female-only rates, bottom panel: male-only rates). ASR is the proportion of the adult population that is male, thus changes in female-biased parameters have a negative effect on ASR and consequently their LTRE statistics are negative. Notation: *h* = mating system index (Eq. 6), *ρ* = hatching sex ratio, Juvenile = sex-biased apparent survival of juveniles, Adult = sex-biased apparent survival of adults.

**Supplementary Figure 2.**
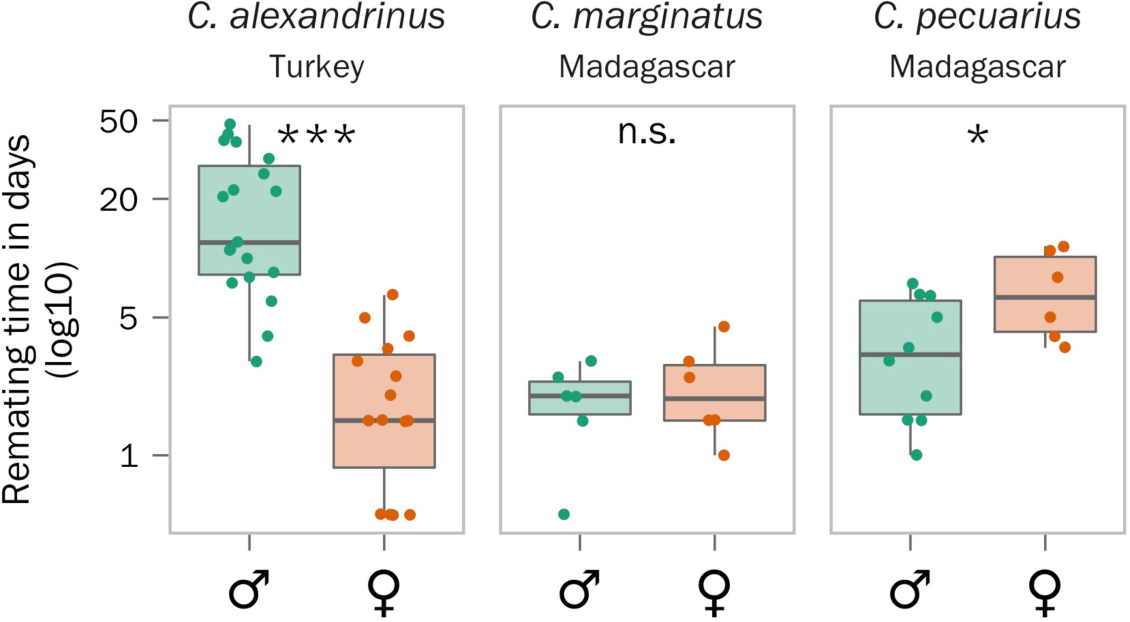
Interspecific variation in sex-specific mating opportunities among three plover species. Experimental assessment of sex-differences in remating times in three of the six populations analysed indicate that *C. alexandrinus* males in Tuzla, Turkey (*n* = 19) take longer to find a mate than females (*n* = 15) after induced divorce. This trend is reversed in *C. pecuarius* (*n*_♂_ = 10, *n*_♀_ = 6) whereas there are no differences in the *C. marginatus* (*n*_♂_ = 6, *n*_♀_ = 6). Significant sex-differences are indicated by asterisks (***: *P* < 0.001, *: *P* < 0.05, n.s.: *P* > 0.05). Figure adapted from Parra et al. (ref. 23).

**Supplementary Figure 3.**
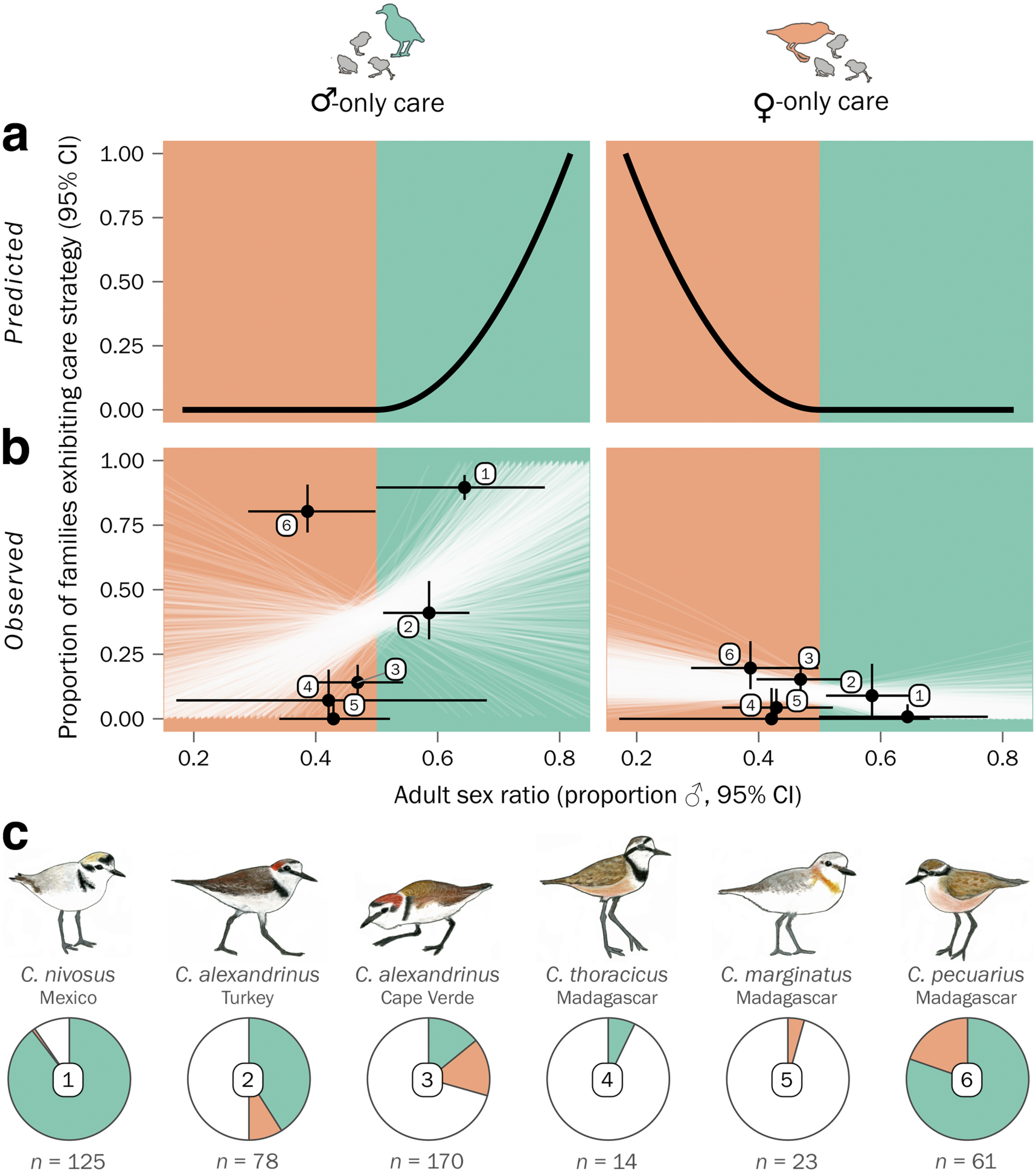
Relationship between uni-parental care and the adult sex ratio. (**a**) Predicted prevalence of male-only care (left panel) or female-only care (right panel) in response to adult sex ratio variation. (**b**) Observed relationship between parental care strategies and adult sex ratio estimates among the six studied populations. Faint white lines illustrate each iteration of the bootstrap, which randomly sampled an adult sex ratio and parental care estimate from each population’s uncertainty distribution and fitted them to the *a priori* exponential model (Eq. 12). (**c**) Proportion of monitored plover families that exhibit parental cooperation (white) or uni-parental care by males (green) or females (orange). Sample sizes reflect the number of families monitored per population, circled numbers correspond to the data point labels shown in panel b.

**Supplementary Figure 4.**
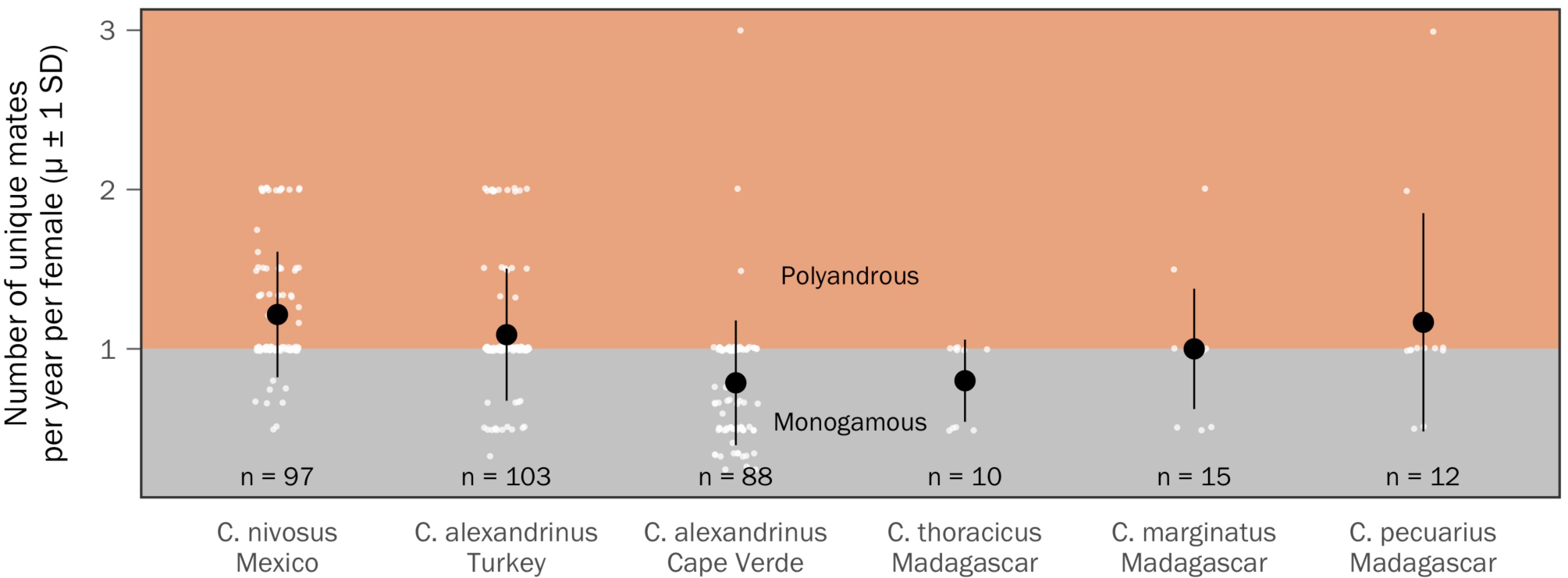
Variation in annual female mating rates (μ) among the six plover populations. Sample sizes indicate the number of individual females in each population that had at least two recorded breeding attempts with identified male(s) during the study. Values below one represent females that bred over multiple years with the same mate (i.e. between season monogamy), whereas values greater that one represent females that have had more than one mate per year (i.e. within season polyandry). Values equal to one represent individuals that have had one mate per year, but have switched mates between years (i.e. between season polyandry but within season monogamy). White data points illustrate individual females, and black points are population averages (*μ* ± 1 SD).

**Supplementary Figure 5.**
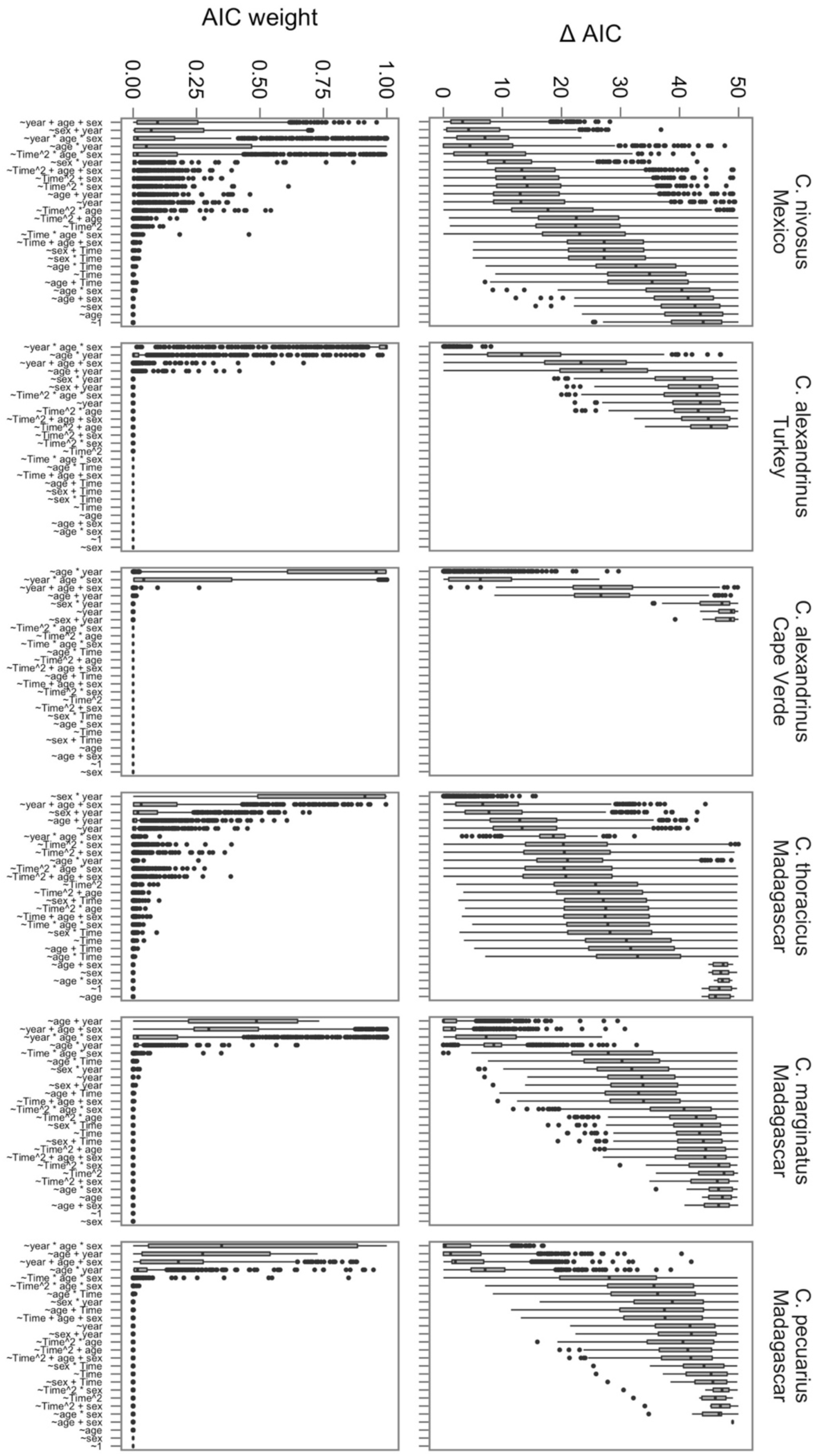
Summary statistics of bootstrapped mark-recapture modelling of juvenile and adult encounter probability. Left panels illustrate variation in AIC_C_ *w_i_*. Right panels illustrate variation in ΔAIC_C_. Model structure of encounter probability (*p*) is shown as labels on the y-axes. See *Methods* for further details.

Supplementary Movie 1

Field methods for capturing and monitoring plovers (11min 46sec): https://drive.google.com/file/d/0B4RO-u-356yiRjNSQ0RUMWpxYlU/view?usp=sharing

## References

1. Arbuthnott, J. An argument for divine providence, taken from the constant regularity observ'd in the births of both sexes. Phil. Trans. 27, 186–190 (1710).

2. Darwin, C. The descent of man and selection in relation to sex. Murray (1871).

3. Donald, P. F. Adult sex ratios in wild bird populations. Ibis 149, 671–692 (2007).

4. Wedekind, C. et al. Persistent unequal sex ratio in a population of grayling (salmonidae) and possible role of temperature increase. Conserv. Biol. 27, 229–234 (2013).

5. Bókony, V., Kövér, S., Nemesházi, E., Liker, A. & Székely, T. Climate-driven shifts in adult sex ratios via sex reversals: the type of sex determination matters. Phil. Trans. 10.1098/rstb.2016.0325 (2017).

6. Petry, W. K. et al. Sex-specific responses to climate change in plants alter population sex ratio and performance. Science 353, 69–71 (2016).

7. Liker, A., Freckleton, R. P. & Székely, T. Divorce and infidelity are associated with skewed adult sex ratios in birds. Curr. Biol. 24, 880–884 (2014).

8. Remeš, V., Freckleton, R. P., Tökölyi, J., Liker, A. & Székely, T. The evolution of parental cooperation in birds. P. Natl. Acad. Sci. USA 112, 13603–13608 (2015).

9. Schacht, R., Rauch, K. L. & Borgerhoff Mulder, M. Too many men: the violence problem? Trends Ecol. Evol. 29, 214–222 (2014).

10. Adimora, A. A. & Schoenbach, V. J. Social context, sexual networks, and racial disparities in rates of sexually transmitted infections. J. Infect. Dis. 191 Suppl 1, S115–22 (2005).

11. Griskevicius, V. et al. The financial consequences of too many men: sex ratio effects on saving, borrowing, and spending. J. Pers. Soc. Psychol. 102, 69–80 (2012).

12. Székely, T., Weissing, F. J. & Komdeur, J. Adult sex ratio variation: implications for breeding system evolution. J. Evolution. Biol. 27, 1500–1512 (2014).

13. Fisher, R. A. The genetical theory of natural selection. Oxford University Press (1930).

14. Trivers, R. L. & Willard, D. E. Natural selection of parental ability to vary the sex ratio of offspring. Science 179, 90–92 (1973).

15. Booksmythe, I., Mautz, B., Davis, J., Nakagawa, S. & Jennions, M. D. Facultative adjustment of the offspring sex ratio and male attractiveness: a systematic review and meta-analysis. Biol. Rev. 92, 108–134 (2017).

16. Székely, T., Thomas, G. H. & Cuthill, I. C. Sexual conflict, ecology, and breeding systems in shorebirds. BioScience 56, 801–808 (2006).

17. Carmona-Isunza, M. C., Küpper, C., Serrano-Meneses, M. A. & Székely, T. Courtship behavior differs between monogamous and polygamous plovers. Behav. Ecol. Sociobiol. 69, 2035–2042 (2015).

18. Veran, S. & Beissinger, S. R. Demographic origins of skewed operational and adult sex ratios: perturbation analyses of two-sex models. Ecol. Lett. 12, 129–143 (2009).

19. Lebreton, J. D., Burnham, K. P., Clobert, J. & Anderson, D. R. Modeling survival and testing biological hypotheses using marked animals: a unified approach with case studies. Ecol. Monogr. 62, 67–118 (1992).

20. Caswell, H. Matrix population models: construction, analysis, and interpretation. Sinauer Associates, Inc. (2001).

21. Emlen, S. T. & Oring, L. W. Ecology, sexual selection, and the evolution of mating systems. Science 197, 215–223 (1977).

22. McNamara, J. M., Székely, T., Webb, J. N. & Houston, A. I. A dynamic game-theoretic model of parental care. J. Theor. Biol. 205, 605–623 (2000).

23. Parra, J. E., Beltrán, M., Zefania, S., Remedios, dos, N. & Székely, T. Experimental assessment of mating opportunities in three shorebird species. Anim. Behav. 90, 83–90 (2014).

24. Amat, J. A., Visser, G. H., Hurtado, A. P. & Arroyo, G. M. Brood desertion by female shorebirds: a test of the differential parental capacity hypothesis on Kentish plovers. P. Roy. Soc. B-Biol. Sci. 267, 2171–2176 (2000).

25. Fromhage, L. & Jennions, M. D. Coevolution of parental investment and sexually selected traits drives sex-role divergence. Nat. Commun. 7, 12517 (2016).

26. Clarke, A. L., Sæther, B.-E. & Røskaft, E. Sex biases in avian dispersal: A Reappraisal. Oikos 79, 429–438 (1997).

27. Küpper, C. et al. High gene flow on a continental scale in the polyandrous Kentish plover Charadrius alexandrinus. Mol. Ecol. 21, 5864–5879 (2012).

28. Eberhart-Phillips, L. J. et al. Contrasting genetic diversity and population structure among three sympatric Madagascan shorebirds: parallels with rarity, endemism, and dispersal. Ecol. Evol. 5, 997–1010 (2015).

29. Eberhart-Phillips, L. J. et al. Sex-specific early survival drives adult sex ratio bias in snowy plovers and impacts mating system and population growth. P. Natl. Acad. Sci. USA 114, E5474–E5481 (2017).

30. Stenzel, L. E. et al. Male-skewed adult sex ratio, survival, mating opportunity and annual productivity in the snowy plover Charadrius alexandrinus. Ibis 153, 312–322 (2011).

31. Küpper, C. et al. Heterozygosity-fitness correlations of conserved microsatellite markers in Kentish plovers Charadrius alexandrinus. Mol. Ecol. 19, 5172–5185 (2010).

32. Kanfi, Y. et al. The sirtuin SIRT6 regulates lifespan in male mice. Nature 483, 218–221 (2012).

33. Clutton-Brock, T. H., Albon, S. D. & Guinness, F. E. Parental investment and sex differences in juvenile mortality in birds and mammals. Nature 313, 131–133 (1985).

34. dos Remedios, N., Székely, T., Küpper, C., Lee, P. L. M. & Kosztolányi, A. Ontogenic differences in sexual size dimorphism across four plover populations. Ibis 157, 590–600 (2015).

35. Zefania, S. et al. Cryptic sexual size dimorphism in Malagasy plovers Charadrius spp. Ostrich 81, 173–178 (2010).

36. Székely, T., Liker, A., Freckleton, R. P., Fichtel, C. & Kappeler, P. M. Sex-biased survival predicts adult sex ratio variation in wild birds. P. Roy. Soc. B-Biol. Sci. 281, 20140342 (2014).

37. Kokko, H. & Jennions, M. Parental investment, sexual selection and sex ratios. J. Evolution. Biol. 21, 919–948 (2008).

38. Hall, L. K. & Cavitt, J. F. Comparative study of trapping methods for ground-nesting shorebirds. Waterbirds 35, 342–346 (2012).

39. Dawson, D. A. et al. Gene order and recombination rate in homologous chromosome regions of the chicken and a passerine bird. Mol. Biol. Evol. 24, 1537–1552 (2007).

40. Küpper, C. et al. Characterization of 36 polymorphic microsatellite loci in the Kentish plover (*Charadrius alexandrinus*) including two sex-linked loci and their amplification in four other Charadrius species. Mol. Ecol. Notes 7, 35–39 (2007).

41. Székely, T. & Cuthill, I. C. Trade-off between mating opportunities and parental care: brood desertion by female Kentish plovers. P. Roy. Soc. B-Biol. Sci. 267, 2087–2092 (2000).

42. Sison, C. P. & Glaz, J. Simultaneous confidence intervals and sample size determination for multinomial proportions. J. Am. Stat. Assoc. 90, 366–369 (1995).

43. Laake, J. RMark: an R interface for analysis of capture-recapture data with MARK. AFSC Processed Report (2013).

44. White, G. C. & Burnham, K. P. Program MARK: survival estimation from populations of marked animals. Bird Study 46, S120–S139 (1999).

45. Bates, D., Mächler, M., Bolker, B. & Walker, S. Fitting linear mixed-effects models using lme4. J. Stat. Softw. 67, 1–48 (2015).

46. Miller, T. E. X. & Inouye, B. D. Confronting two-sex demographic models with data. Ecology 92, 2141–2151 (2011).

47. Caswell, H. Perturbation analysis of nonlinear matrix population models. Demogr. Res. 18, 59–116 (2008).

48. Carmona-Isunza, M. C. et al. Adult sex ratio and operational sex ratio exhibit different temporal dynamics in the wild. Behav. Ecol. 28, 523–532 (2017).

49. Jenouvrier, S., Caswell, H., Barbraud, C. & Weimerskirch, H. Mating behavior, population growth, and the operational sex ratio: a periodic two-sex model approach. Am. Nat. 175, 739–752 (2010).

50. R Core Team. R: a language and environment for statistical computing. (2017). doi:ISBN 3-900051-07-0

